# Determinants of Transcription Initiation Directionality in Metazoans

**DOI:** 10.1101/224642

**Authors:** Mahmoud M. Ibrahim, Aslihan Karabacak, Alexander Glahs, Ena Kolundzic, Antje Hirsekorn, Alexa Carda, Baris Tursun, Robert P. Zinzen, Scott A. Lacadie, Uwe Ohler

## Abstract

Divergent transcription from promoters and enhancers is pervasive in many species, but it remains unclear if it is a general and passive feature of all eukaryotic cis regulatory elements. To address this, we define promoters and enhancers in *C. elegans, D. melanogaster* and *H. sapiens* using ATAC-Seq and investigate the determinants of their transcription initiation directionalities by analyzing genome-wide nascent, cap-selected, polymerase run-on assays. All three species initiate divergent transcription from separate core promoter sequences. Sequence asymmetry downstream of forward and reverse initiation sites, known to be important for termination and stability in *H. sapiens*, is unique in each species. Chromatin states of divergent promoters are not entirely conserved, but in all three species, the levels of histone modifications on the +1 nucleosome are independent from those on the -1 nucleosome, arguing for independent initiation events. This is supported by an integrative model of H3K4me3 levels and core promoter sequence that is highly predictive of promoter directionality and of two types of promoters: those with balanced initiation directionality and those with skewed directionality. Lastly, *D. melanogaster* enhancers display variation in chromatin architecture depending on enhancer location, and *D. melanogaster* promoter regions with dual enhancer/promoter potential are enriched for divergent transcription. Our results point to a high degree of variation in regulatory element transcription initiation directionality within and between metazoans, and to non-passive regulatory mechanisms of transcription initiation directionality in those species.

---

The application of deep sequencing assays led to the unanticipated observation that the promoters of many genes are transcribed in both directions, a phenomenon dubbed divergent transcription. In divergent promoters, transcripts made in the direction antisense to the annotated gene are non-protein-coding and highly unstable such that they can typically only be detected in assays enriching for nascent RNA. Divergent transcription is pervasive across many eukaryotes including yeast, *C. elegans, M. musculus* and *H. sapiens*^1–5^, though is highly depleted in *D. melanogaster*^6^.

In mammals, the asymmetric output of divergent promoters was suggested to be the result of a post-transcriptional competition model between the splicing machinery and the cleavage/polyadenylation machinery such that enriched splice site sequences lead to transcript extension and stabilization in the forward direction, whereas enriched cleavage sequences lead to transcription termination and RNA degradation by the nuclear exosome complex in the reverse unstable direction^7,8^. A different, Nrd1-complex mediated mechanism was found to destabilize divergent promoter transcripts in yeast^5,9,10^.

These observations are unable to fully explain transcription directionality since considerable variation in forward/reverse transcription rates was measured by nascent RNA data^4,6,11^. Divergent promoters initiate transcription from two separate core promoters upstream antisense to each other within a single nucleosome depleted region (NDR), forming two distinct polymerase pre-initiation complexes (PICs)^11–14^. Differences in the sequence-encoded strengths of the forward- and reverse-directed core promoters were suggested to drive variation in promoter directionality in *H. sapiens* HeLa cells^11,15^. Therefore, asymmetric output of mammalian divergent promoters is potentially sequence-encoded at both transcription initiation and post-transcriptional termination/degradation.

The level of divergent transcription is also reflected in a unique promoter chromatin environment exemplified primarily by differences in levels and distribution of methylation on lysine 4 of histone H3 (H3K4me1/2/3) upstream of the promoter NDR^11,16^. H3K4 methylation and other histone post-translational modifications (PTMs) on promoter NDR-flanking nucleosomes are known to influence transcription initiation and elongation rates via direct physical interactions with PICs^17–19^, which may contribute to directional variation of transcription initiation within promoter NDRs.

Divergent transcription is also observed in distal gene regulatory elements such as enhancers, i.e. with unstable-unstable transcript pairs, and has been recently regarded as a defining feature of active enhancers in mammals^12,20,21^. While enhancers have been long known to feature different chromatin states than those of promoters^22^, recent studies have led to the hypothesis that promoters and enhancers are not distinct types of regulatory elements since they both feature divergent transcription, with H3K4 methylation states varying according to differences in transcription initiation rates^12,23^. Of note, the striking similarities in architecture between promoters and enhancers does not necessarily translate to functional equivalence^24^.

While divergent transcription in mammals is reflected in both DNA sequence and chromatin, the precise contribution of sequence and chromatin features to transcription initiation directionality is not well understood. To reconcile seemingly contradictory observations about the prevalence of divergent transcription in different eukaryotes, as well as the mechanisms enforcing it, we quantify the directional relationships between promoter sequence, histone PTMs, and transcription initiation for *Drosophila melanogaster, Caenorhabditis elegans*, and *Homo sapiens*. We observe strict directional correlations between core promoter sequence strengths and initiation quantity in all three species, as well as highly directional correlations between active histone modifications upstream and downstream of promoter NDRs in all three species. We find forward/reverse histone modification levels and core promoter sequence strengths alone to be highly predictive of promoter initiation directionality and to suggest two, potentially mechanistically distinct, promoter types. Sequence content asymmetry adjacent to promoter NDRs is distinct across species and suggests species-specific mechanisms for post-transcriptional contributions to transcript directionality. Finally, low-level divergent transcription initiation is detected from active enhancers in all three species, with putative enhancer activity strongly enriching for divergent transcription in *D. melanogaster* promoters.

## Results

### Sequence features differentially contribute to promoter NDR directionality across species

We performed the assay for transposase-accessible chromatin (ATAC-seq) on *D. melanogaster* S2 cells and C. *elegans* whole L3-stage to compare with previously published data in the *H. sapiens* cell line GM12878^25^. NDRs were computed using peak-calling with the JAMM algorithm^26^ on all three datasets and the resulting peaks were annotated as promoters based on proximity to an annotated Transcription Start Sites (TSS, see Methods). This yielded 18067 promoter NDRs in the *H. sapiens* cell line, 6926 in the *D. melanogaster* cell line, and 10912 in the L3-stage whole *C. elegans*.

To assess directionality of transcription initiation for the detected NDRs, we used previously published G/PRO-cap datasets in *H. sapiens* GM12878, *D. melanogaster* S2 cells and L3-stage whole *C. elegans*^4,12,27^. Counting read starts from these nascent TSS datasets within promoter NDRs resulted in 14371 (79%), 6280 (91%), and 10786 (99%) regions in *H. sapiens, D. melanogaster*, and *C. elegans*, respectively, with at least one TSS read on at least one strand. Evaluating the forward (annotated gene) nascent TSS read counts against those on the reverse strand for these groups produces a minimally biased view of promoter transcription initiation directionality across the three species (Figure 1A). Focusing beyond basal, likely inactive subpopulations (lower left corners), *H. sapiens* GM12878 cells show some correlation between forward and reverse signal, but with a substantial skew toward the x-axis, reflecting a larger number of promoters with biased directionality toward the annotated gene. Previously published peak calls from DNasel-seq and 5’-GRO-seq TSS data in HeLa cells also show bias in directionality toward annotated genes (Figure S1A), consistent with published observations^11^. The skew of initiation toward annotated genes is exacerbated in *D. melanogaster*, with most data points lying tight along the x-axis (Figure 1A), whereas *C. elegans* shows a distribution in between *D. melanogaster* and *H. sapiens*, consistent with meta analyses (Figure 1A) and with previous reports^4,6^.

**Figure 1.**
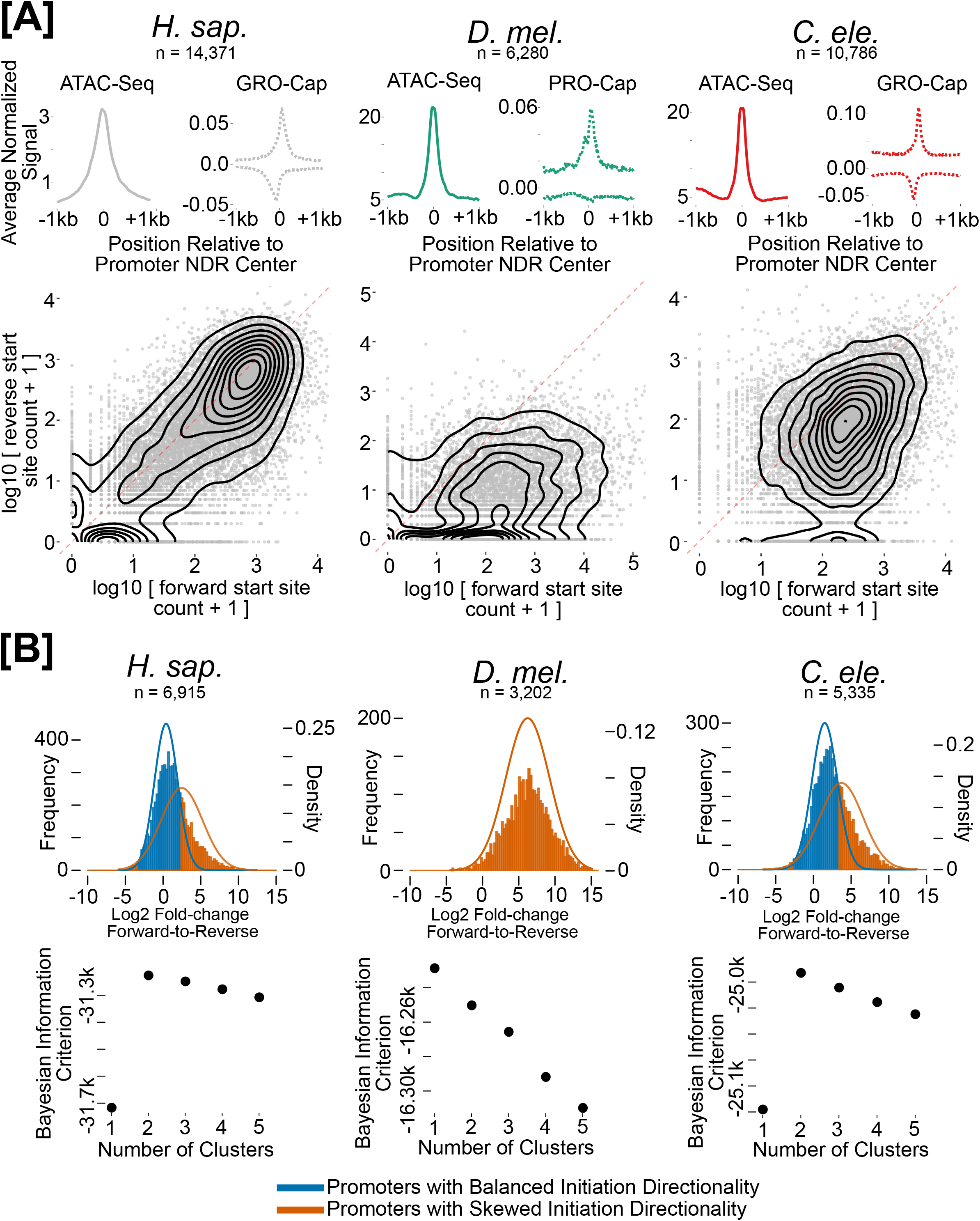
Variation of transcription directionality within and across species. **A**. Meta analyses for average depth-normalized ATAC-seq (solid line) coverage and zero-to-one-scaled PRO/GRO-cap (dotted line) coverage relative to promoter NDR midpoints defined by ATAC-seq are displayed on top for each species. Forward direction (annotated gene) versus reverse direction cap-selected PRO/GRO-seq counts displayed as contour scatter plots for the same promoter NDRs in *H. sapiens* GM12878 (left) *D. melanogaster* S2 cells (middle) and whole L3 *C. elegans* (right). **B**. Mixture models (top) and BIC cluster analysis (rbottom) of forward/reverse cap-selected GRO/PRO-seq count ratios for promoter NDRs containing significant forward initiation. A pseudo count of 1 was added to numerators and denominators. Lines represent density of theoretical distributions learned, histograms represent observed ratios.

To estimate directionality subpopulations, we applied Gaussian mixture modeling to log forward-to-reverse ratios in promoters that showed substantial expression in the forward direction (see Methods). Bayesian information criteria analysis of cluster numbers suggested 2, 1, and 2 mixture components as optimal for *H. sapiens, D. melanogaster*, and *C. elegans*, respectively. These analyses clearly show that promoter architecture is diverse across species with respect to the directionality of transcription, with *D. melanogaster* data in particular supporting only one type of promoter with little divergent transcription, 29% promoter NDRs with initiation highly skewed toward the annotated gene for *H. sapiens* GM12878, and 34% for stage L3 *C. elegans* (Figure 1B).

We then selected high confidence divergent promoters where both the forward, annotated-gene side, and the reverse, un-annotated side of the NDR initiate transcription above a stringent background model (see Methods). Since only 441 promoter NDRs met these criteria in S2 cells, we wondered whether this group reflects true divergent promoters. We generated PEAT data in S2 cells^28^, an assay which measures TSSs of stable polyadenylated transcripts and found the reverse signal of the selected divergent promoters to be preferentially depleted in stable transcripts (Figure S1B). Therefore, albeit much less frequent than in *H. sapiens* or *C. elegans*, the selected *D. melanogaster* group is likely to correspond to true divergent promoters.

The initiation pattern (i.e. the distribution of start site read counts across positions within a promoter; Figure 2A) has been shown to correlate with other promoter properties such as core promoter sequence elements and expression level^29–31^ so we first measured these distributions, using a previously described entropy-based metric^32^, for the forward and reverse TSSs within divergent promoters. Overall, there is a high degree of overlap between the distributions of forward and reverse initiation pattern scores in all three species, suggesting that reverse TSSs in our stringent groups are not randomly located events (Figure S1C). To quantify the role of core promoter sequence in directing reverse initiation from divergent promoter NDRs, we turned to a previously developed position-specific Markov chain model of TSS sequences^33^ and applied each species-specific model to the forward and reverse TSS sequences from the stringently-selected divergent NDR groups (see Methods; Figure 2A). As we previously reported for *H. sapiens* HeLa cells^11^, reverse TSSs from GM12878, *D. melanogaster* S2, and *C. elegans* L3 all score well compared to random controls taken from the center of the divergent NDRs (Figure 2B). Together with the presence of well positioned TATA and initiator consensus motifs around reverse TSSs (Figure S1D), these data strongly suggest that all three species initiate reverse-directed transcription from reverse-directed core promoter sequences within NDRs. Sequences at forward initiation sites for *D. melanogaster* show substantially increased model scores compared to *H. sapiens* and *C. elegans* core promoters, while the *D. melanogaster* forward score distribution is more separable from the *D. melanogaster* reverse distribution, suggesting that positional and directional sequences within core promoters are highly prevalent in *D. melanogaster*, consistent with previously reported observations^34,35^, and that this may contribute to the overall scarcity of divergent promoters in *D. melanogaster* (Figure 1A).

**Figure 2.**
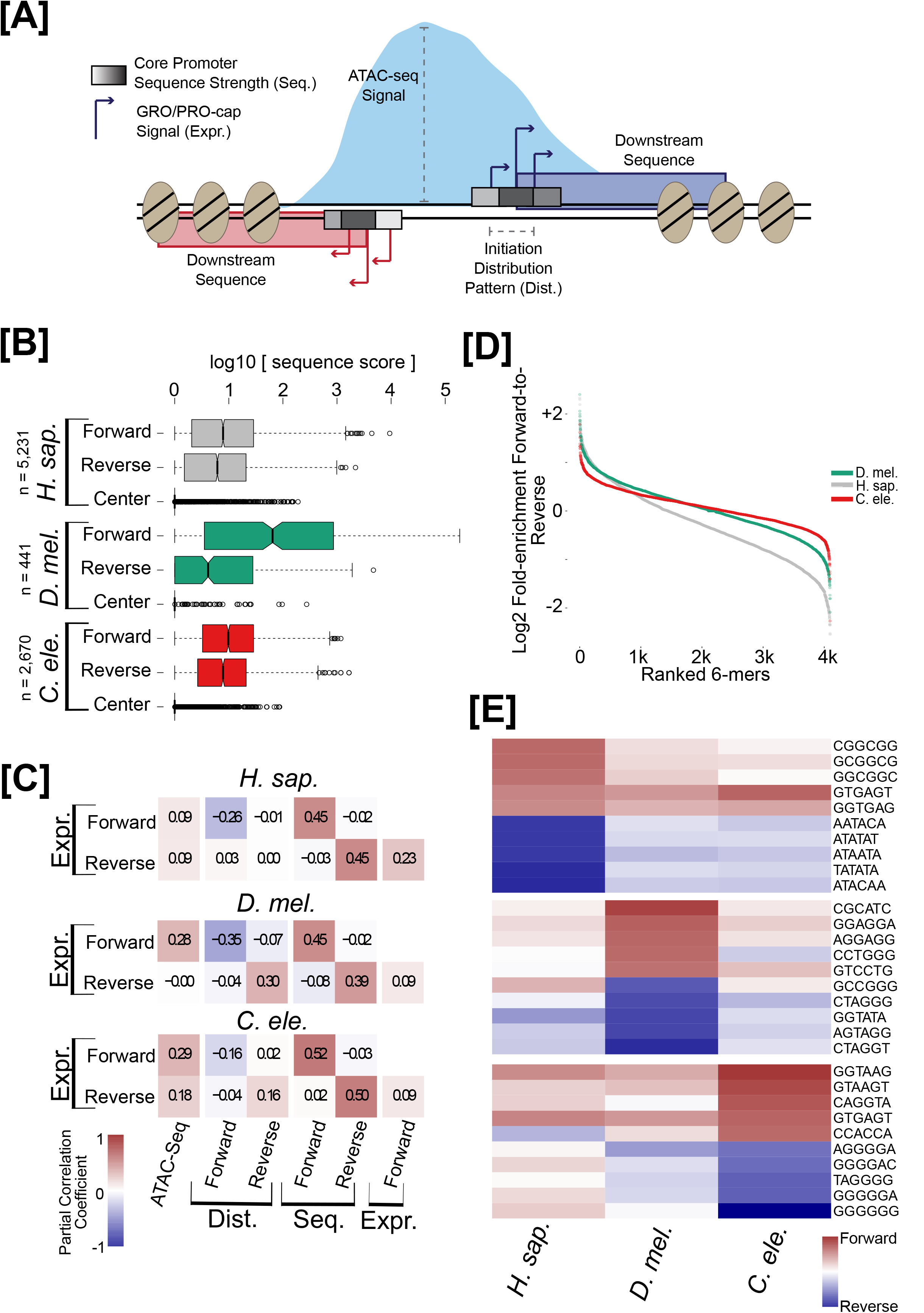
Asymmetric sequence features contribute to variation of transcription directionality within and across species. **A**. Schematic of features measured for stringently-selected divergent promoters. **B**. Core promoter sequence model scores at significant forward and reverse TSS modes for promoter NDRs in all three species (see Methods). Center positions between forward and reverse TSSs serve as negative controls. **C**. Partial correlation analysis between total cap-selected PRO/GRO-seq counts (Expr.), ATAC-seq signal, TSS distribution entropy (Dist.) and core promoter sequence score sums (Seq.) in forward and reverse directions for promoter NDRs with significant forward and reverse TSSs (see Supplementary Table 1 for full partial correlation table). **D**. All 6-mer sequences ranked by, and plotted against, their count ratios from 500 bp windows downstream of forward and reverse TSSs for the same divergent promoter NDRs. **E**. Top-5 (red) or bottom-5 (blue) 6-mers (see **D**.) from each species with their respective scaled forward/reverse count ratios in each other species.

To determine pairwise relationships between sequence content and transcription initiation features, we turned to rank-based partial-correlation analysis, which examines pairwise correlations between each two features removing confounding effects due to all the other features. We applied the partial-correlation analysis to ATAC-seq counts, forward and reverse initiation rates as measured by cap-selected G/PRO-seq, initiation distribution entropy scores, and core promoter sequence model scores of divergent promoter NDRs in each species (Figure 2A; Figure 2C). Strikingly, all three species show correlations between core promoter sequence model scores for forward and reverse TSSs with their respective initiation rate counts, but forward model scores do not relate to reverse initiation counts and reverse model scores do not relate to forward initiation counts. These observations confirm the key contribution of reverse-directed core promoter sequences to divergent transcription from promoter NDRs.

Asymmetric sequence content downstream of forward and reverse TSSs has been shown to influence transcription elongation directionality^7,8^. To compare these asymmetries across organisms, we adopted the approach taken by Almada et al., wherein the ratio of forward to reverse counts for all six-mer sequences is calculated (Figure 2A; Figure 2D)^7^. Comparing the 5 most forward- and reverse-enriched sequences found for each species shows that these asymmetries are quite unique to each organism: these six-mers are only enriched in one species, with the 5’ splice site consensus GTGAGT as only exception (Figure 2E). The observations for *H. sapiens* are consistent with previous reports of high enrichment in the forward direction for the consensus 5’ splice site sequence and high enrichment in the reverse direction for AT-rich, cleavage-like sequences. The 5’ splice site is also forward-enriched in *C. elegans* and somewhat so in *D. melanogaster*, but neither *D. melanogaster* nor *C. elegans* shows top reverse enrichment of AT-rich six-mers; something that is also reflected in average positional GC content (Figure S1E). The most highly reverse-enriched six-mers in *C. elegans* contain G stretches which is also reflected in a striking pattern of average positional GC-skew (Figure S1E). Together these observations suggest that sequence asymmetry downstream of forward and reverse TSSs of divergent promoter NDRs is quite distinct across species; together with the known Nrd1-mediated model in yeast^5,9,10^, this suggests that the splicing/cleavage competition model of transcript elongation may not apply outside of vertebrates.

### Promoter chromatin environment is directional and varies across species

We took advantage of the high resolution of ATAC-Seq and PRO/GRO-cap assays to ask whether differences exist between the chromatin organization of *H. sapiens, D. melanogaster* and *C. elegans* promoters. Indeed, we found *D. melanogaster* and *C. elegans* promoter NDRs to be significantly smaller on average than *H. sapiens* promoter NDRs (Figure S2A) and their transcription initiation sites to be closer to the +1 nucleosome than in *H. sapiens* (Figure S2B). We generated ChIP-seq data for H3K4me2 in *D. melanogaster* S2 cells to combine with publicly available datasets for H3K4me1, H3K4me3, and H3K27ac^36^, as well as all four modifications in *C. elegans* and *H. sapiens*^37,38^ (see Methods). We then characterized combinatorial states for those four PTMs for each organism independently at 10-bp resolution, using a multivariate Gaussian Hidden Markov Model (see Methods). We detected 11 chromatin states, which we named according to their positional trends: three promoter states (P1, P2, and P3) characterized by H3K27ac and H3K4me3 with different H3K4me1 and H3K4me2 levels, two transcription elongation states (EL1 and EL2) containing H3K4 methylation without H3K27ac, two enhancer states (E1 and E2) with H3K27ac and H3K4me1 or H3K4me2 but not H3K4me3, two H3K4me1-only states, one H3K4me2-only state, and a background state (Figure 3A).

**Figure 3.**
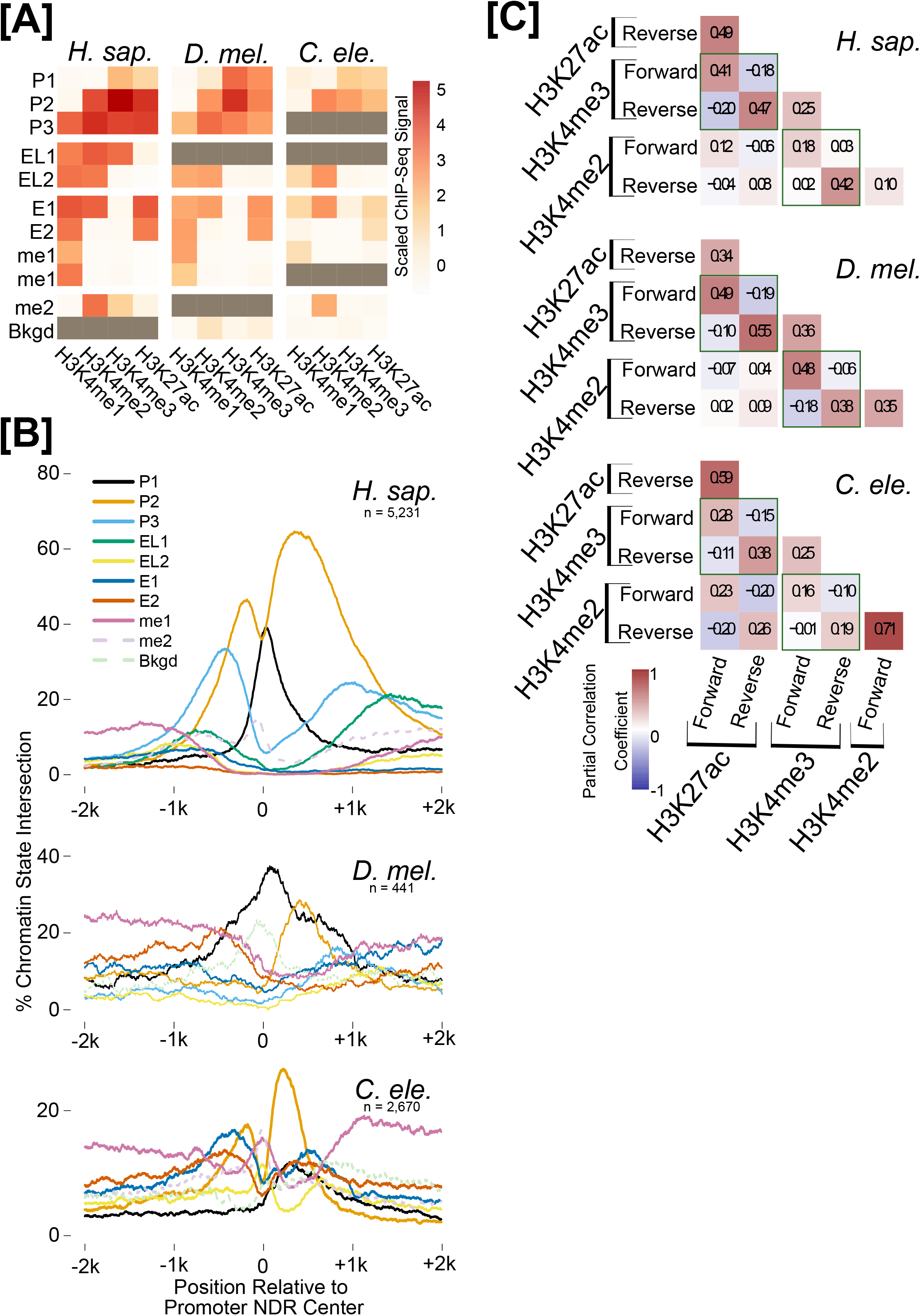
Promoter histone PTM states are strictly directional. **A**. PTM signal heatmap for measured HMM states in all three species, showing the mean signal in each state learned by the model. (gray boxes indicate the state was not detected in the respective organism) **B**. Histone PTM state positional coverage for promoters with significant forward and reverse TSSs. **C**. Partial correlation analysis between active promoter histone PTMs in forward and reverse directions for the same promoter NDRs (see Supplementary Table 2 for full partial correlation table).

Consistent with previous observations^39^, the discovered chromatin states are similar across species, but there are interesting differences in their spatial arrangement around promoter NDRs. The forward direction of divergent NDRs in *H. sapiens* and *D. melanogaster* show a similar cascade of P1-P2-P3 reminiscent of our previous observations in HeLa cells (Figure 3B)^11^, while the forward direction in the *C. elegans* divergent NDRs is dominated by P2, reflecting the relative confinement of H3K27ac, H3K4me2, and H3K4me3 to the +1 nucleosome (Figure 3B; Figure S2C). In the reverse direction, *H. sapiens* and *C. elegans* are enriched for P2 similar to our findings in HeLa cells^11^, whereas *D. melanogaster* divergent promoters display an enrichment of P1 on their -1 nucleosome, a state that is enriched only in the forward direction for *H. sapiens* and *C. elegans* (Figure 3B, compare black lines for P1 and orange lines for P2). Therefore, the histone PTM spatial distribution and combinations, upstream and downstream of divergently transcribed promoter NDRs, vary from species to species.

We then performed partial-correlation analysis as before (see Figure 2C), using the maximum ATAC-Seq signal in the NDR and the maximum forward and reverse levels of histone modifications in a 1kb window downstream and upstream of the promoter NDR (Figure 3C). A strictly directional correlation can be seen between H3K27ac and H3K4me3, such that positive relationships are observed between the PTM levels both in the forward and the reverse directions, but forward levels show inverse correlations with reverse levels (Figure 3C, follow green squares). Positive correlations are seen between the same PTM on the forward and reverse sides of divergent promoter NDRs in all three species, but this is an expected confounding factor due to the low resolution of the ChIP-seq assay relative to the promoter NDR width.

### Sequence and chromatin features are predictive of promoter directionality

We sought to integrate both core promoter sequence and histone PTM in a predictive model of transcription initiation directionality, defined as the ratio of forward to reverse initiation counts. We constructed a model that simultaneously learned a mixture of two linear models, in which different coefficients for two features (core promoter sequence scores and H3K4me3 PTM levels) are assigned for each linear model separately. Therefore, both transcription initiation directionality ratio and promoter type are simultaneously predicted when the model is trained (see Methods).

Assigning each promoter to the type predicted by the model and comparing the distributions of experimentally measured forward/reverse initiation ratios leads to two distributions highly similar to the bimodal directionality distributions (Figure 4A, Figure 1B), indicating that the model is able to discern both the directionally balanced and directionally skewed promoter types based only on core promoter sequence scores and H3K4me3 levels. Comparing the predicted forward/reverse transcription initiation ratio against the experimentally measured value for each promoter leads to a correlation of 0.69, suggesting that these two features together are highly predictive of promoter transcription initiation directionality (Figure 4A, Figure S3A). Furthermore, assessing the full model and two models trained on sequence only and H3K4me3 only shows that core promoter sequence score is the more important feature for the model’s performance overall (Figure 4B). Regression coefficients indicate that core promoter sequence scores appear to be more influential for predicting directionality in promoters with skewed directionality than in directionally balanced promoters (Figure S3B). This analysis lends further support to the hypothesis that promoter directionality is variable and functionally determined by promoter NDR sequence content and adjacent histone PTM levels^11,15,40^.

**Figure 4.**
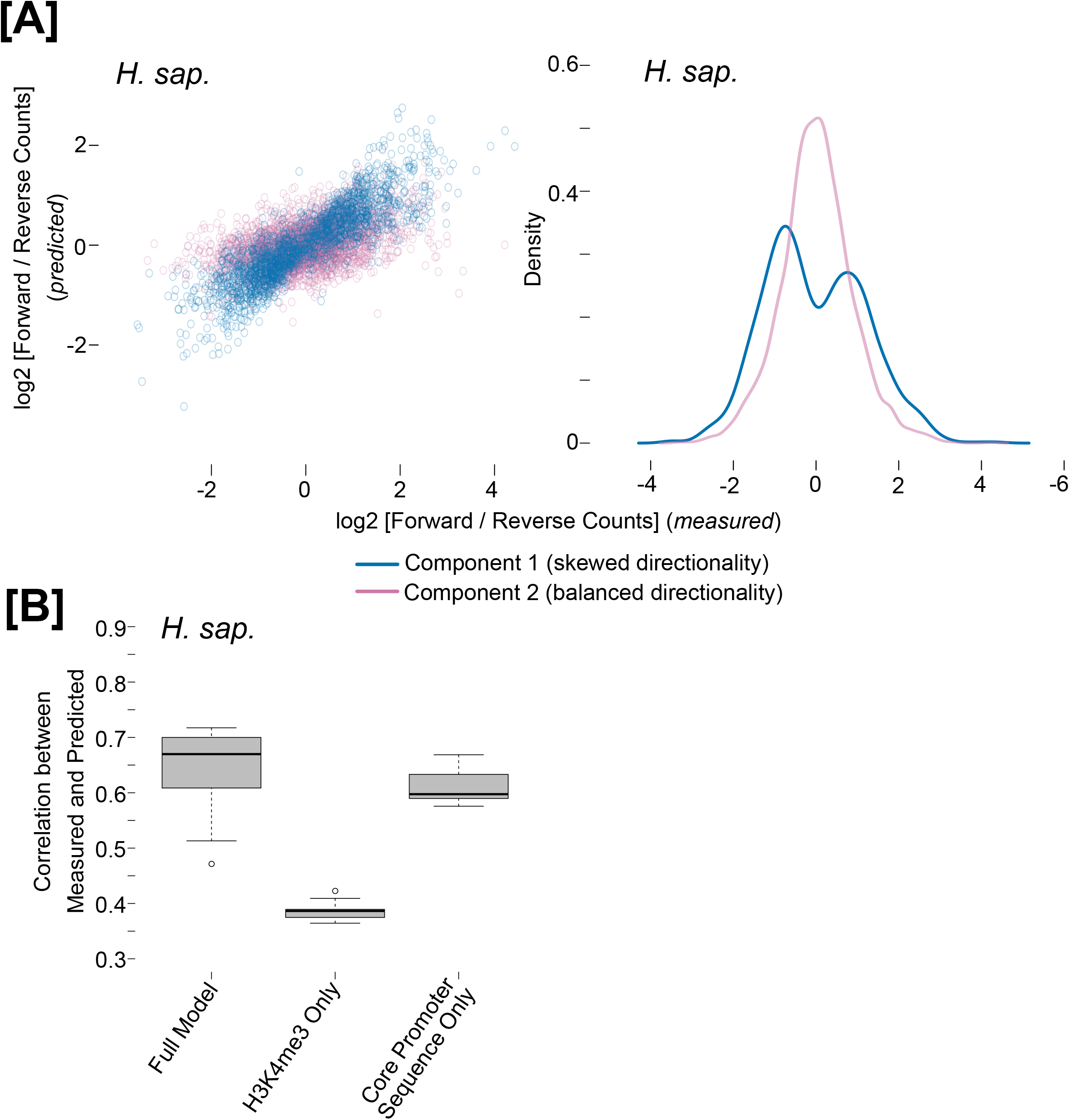
H3K4me3 and core promoter sequence are predictive of promoter directionality. Predicted vs. measured transcription directionality in *H. sapiens* promoter regions using a mixture linear model (left). Density plot of promoter counts for different directionality groups (right). Model is trained on all promoter regions and skewed (blue) and balanced (pink) directionality model components are displayed separately. **B**. 10-fold cross-validation of three models including a model trained using core promoter sequence score ratios and H3K4me3 signal ratios, core promoter sequence score ratios only and H3K4me3 ratios only. Boxplots show the correlation between predicted and measured transcription directionality for the test sets.

### Variation of distal regulatory architecture

To investigate the transcriptional and histone PTM levels of distal enhancer NDRs, we selected ATAC-seq peaks intersecting at least H3K4me1 and H3K27ac from each species and situated far from annotated genes on both strands (ie. potentially active enhancers^41^, see Methods). Forward and reverse nascent transcription initiation counts for those regions indicate that all three species display enhancers with transcripts in both directions (Figure 5A). As expected, distal NDRs from all three species are depleted of promoter and elongation associated states (Figure 3A, Figure 5B, Figure S4A). *D. melanogaster* and *H. sapiens* distal NDRs show a progressive increase in transcription initiation levels as they intersect peaks for H3K4me1 only, H3K4me1/H3K27ac only, and H3K4me1/H3K4me2/H3K27ac only, consistent with current models of enhancer transcription and activation^42^ (Figure S4B), but this was not observed for *C. elegans*. While many distal accessible regions marked by H3K4me1/H3K4me2/H3K27ac harbor cap-selected PRO/GRO-seq signal (Figure S4B), it remains unclear if all active enhancers have these histone marks and/or produce eRNAs.

**Figure 5.**
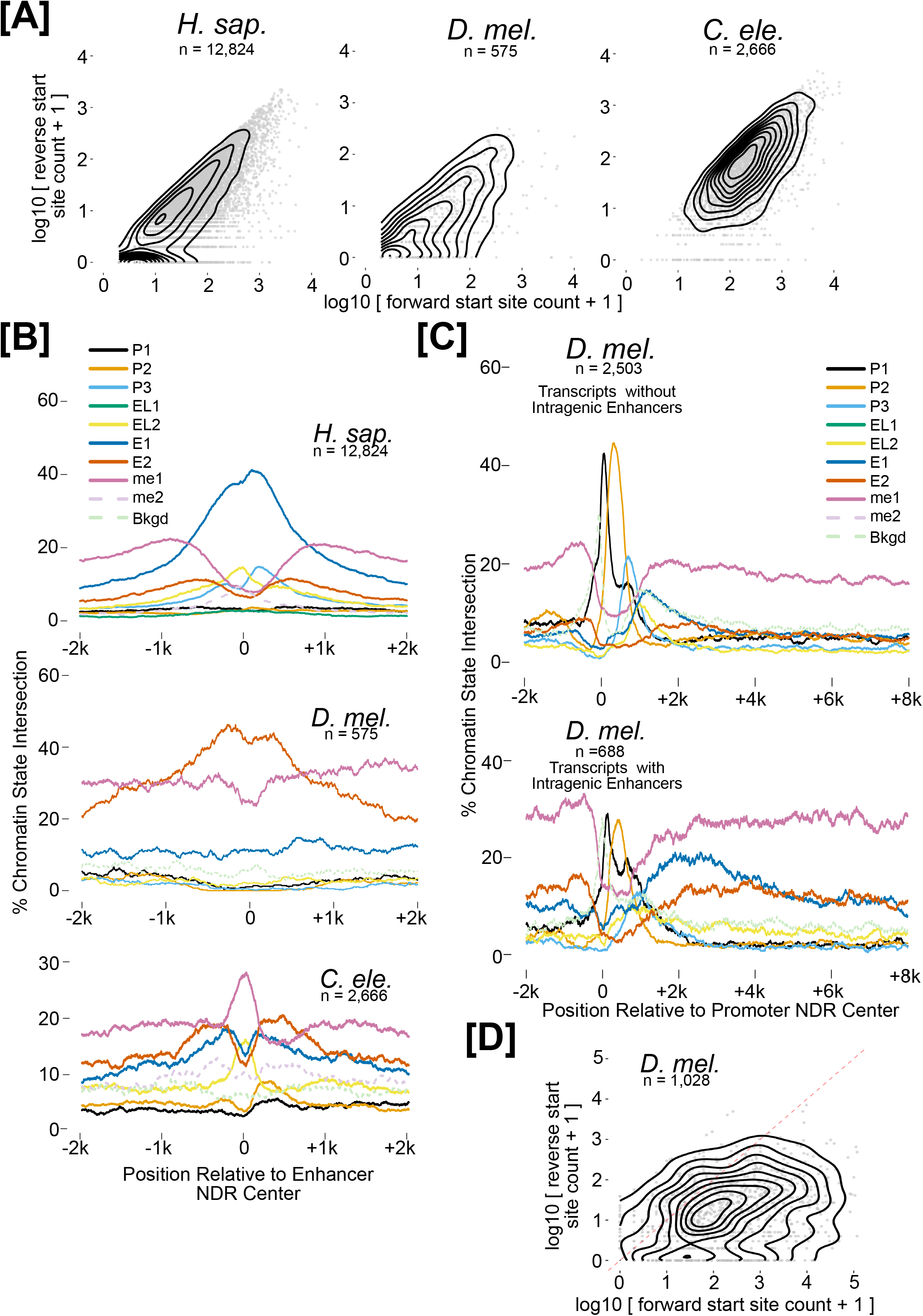
Distal regulatory architecture varies across species. **A**. Forward versus reverse direction cap-selected G/PRO-seq counts for ATAC-seq-defined intergenic NDRs, which intersected at least H3K4me1 and H3K27ac in *H. sapiens* GM12878 (left), *D. melanogaster* S2 cells (middle) and whole L3 *C. elegans* (right). Forward and reverse are the strands with higher and lower counts respectively. **B**. Histone PTM state positional coverage for the same NDRs as in **A. C**. Histone PTM state positional coverage for expressed promoters whose transcripts are either without (top) or with (bottom) intragenic nucleosome-depleted STARR-seq peaks. **D**. Forward versus reverse direction cap-selected PRO-seq counts for promoter NDRs that intersect STARR-seq enhancer peaks.

While *C. elegans* and *H. sapiens* intergenic enhancers show bimodal patterns of both enhancer state 1 (E1; H3K4me1/H3H4me2/H3K27ac) and enhancer state 2 (E2; H3K4me1/H3K27ac), *D. melanogaster* shows relatively low coverage of E1 and H3K4me2, while maintaining a bimodally-enriched pattern for E2 (Figure 5B, Figure S4A). This prompted us to ask if an alternative set of NDRs within gene bodies might contain the E1 state. To address this, we classified gene bodies into those that do and do not contain NDRs intersecting experimentally determined enhancers as defined by the STARR-seq assay^43^. This displays a striking enrichment of both E1 and E2 states well downstream of the promoter region, only for the group containing S2 cell active STARR-seq enhancers (Figure 5C). Together with the state pattern in *D. melanogaster* intergenic NDRs (Figure 5B), these observations suggest that *D. melanogaster* enhancers tend to have different chromatin architecture depending on whether or not they fall within a gene.

Since some *D. melanogaster* promoters were also found to show potential enhancer activity in S2 cells^43^, we specifically selected promoter-annotated NDRs that intersected STARR-seq peaks and detected a strong enrichment for divergent transcription initiation compared to promoter NDRs not intersecting STARR-seq peaks (Figure 5D, Figure S4C). This suggests that divergent transcription might indeed be a strong indicator for enhancer activity, consistent with reporter-based activity assays of divergently transcribed *H. sapiens* enhancers^44^.

## Discussion

We observe strict directional correlations between core promoter sequence strengths and initiation rates in the forward and reverse directions from promoter NDRs. Therefore, forward and reverse directed transcription events are measurably independent from each other, consistent with previous observations of separate preinitiation complex formation and clear separation of initiation sites^11,12,14,45^. Our analysis also indicates a strict directional positive correlation between different histone PTMs on the +1 and −1 nucleosomes of promoter NDRs in all three species (Figure 3C). We and others have previously reported a directional histone PTM arrangement around promoter NDRs likely reflecting differences in initiation directionality^11,40,45^, though RNAPII kinetics and RNAPII PTMs are also likely to contribute^16^. A promoter directionality model emerges whereby directional synergy between core promoter sequences and histone PTMs in the forward and reverse directions determines fitness in a competition for a common pool of RNAPII to initiate transcription at the downstream or upstream edges of promoter NDRs^15^. We tested this idea using a linear regression model and found core promoter sequence strength and H3K4me3 levels to be predictive of transcription initiation directionality.

Using this mixture model, we could also distinguish two separate groups of promoters, which we define more precisely here as promoters with skewed directionality and promoters with balanced directionality. Those two groups show differences in how the synergy between sequence and chromatin is coordinated: while promoters with skewed directionality are mainly determined by core promoter sequence, histone PTMs play a bigger role in determining directionality of balanced promoters. This again poises divergent transcription as a potentially regulatory mechanism, rather than a passive consequence of transcription initiation.

We propose a refined picture of transcriptional directionality in which (a) skewed directionality is enforced at genuine endogenous promoters, where one side acquired functionality to transcribe a relatively more functional trans-acting (m)RNA at relatively higher levels, consistent with recent studies by Jin et al.^46^; (b) transcription of a divergent product (functional or not) may also act as a tuning mechanism for the initiation rates of a functional, oppositely-oriented counterpart; (c) the directional variation of initiation across NDRs is determined by directionally competing sequence and chromatin features; and (d) apparently species-specific mechanisms ensure that any divergent, nonfunctional transcripts are efficiently degraded.

Our finding that enhancer activity overlapping annotated promoter regions in *D. melanogaster* S2 cells enriches for divergent transcription was also shown in *H. sapiens* cells^47^ and is consistent with previous observations using the CAGE assay in mammalian cells^44^, potentially suggesting a function for divergent transcription at enhancers. Transcription initiation may help to position nucleosomes, thereby ensuring accessibility to the DNA by transcription factors, and nascent eRNA may act to compete with chromatin for nucleic acid binding factors/complexes^48^. On the other hand, enhancers have a different functional requirement than promoters: they do not need to produce stable transcripts at possibly high levels, as exemplified by recent studies^24^. It is possible that enhancers with skewed directionality act in a mechanistically distinct way, e.g. as promoters for lncRNAs which subsequently act *in trans* as transcriptional regulators, but such distinctions remain to be addressed. As the nascent transcriptomes of more eukaryotes are profiled, we anticipate that a wide range of transcription directionality tendencies will be observed with different chromatin-sequence synergy mechanisms.

## Methods

### *C. elegans* ATAC-seq

*C. elegans* wild-type strain N2 was grown on OP50 bacteria at 20°C as described before (Brenner, 1974). Embryos were harvested from adults by sodium hypochlorite treatment and grown until third larval instar (L3). Synchronized L3 animals were washed 5 times in M9 buffer and collected on ice. Nuclei were isolated using a glass Dounce homogenizer with 50 strokes tight-fitting insert in buffer A (15 mM Tris-HCl pH7.5, 2 mM MgCl2, 340 mM sucrose, 0.2 mM spermine, 0.5 mM spermidine, 0.5 mM phenylmethanesulfonate [PMSF], 1mM DTT, 0.1% Trition X-100 and 0.25% NP-40 substitute) as described before (Ooi et al., 2010; Steiner and Henikoff, 2014). The debris were removed by spinning at 100×g for 5 min and nuclei were counted by Methylene blue staining. 100.000 nuclei per sample were pelleted by spinning at 1000×g for 10 min and proceeded immediately to transposition step of the ATAC-seq protocol (Buenrostro et al., 2013). Libraries were amplified for a total of 13 or 14 cycles.

### *D. melanogaster* S2 cell ChIP-seq, ATAC-seq, and PEAT

For ChIP-seq, *D. melanogaster* S2 cells were obtained from Life Tech (#R69007) and grown at 25°C in Schneider’s Cell medium (Life Tech, #21720024) with 10% FBS (Sigma, #F7524) and 10% L-Glutamine (Sigma, #G7513) without antibiotics. Cells were grown in T75 flasks at 25°C to a confluency of ~70%. For ATAC-seq, cells were grown at 25°C in ExpressFive SFM medium (Life Tech, #10486025) with 10% heat-inactivated FBS (Life Tech, #16000044) and 12% L-Glutamine (Life Tech, #25030024) and 1% Penicillin-Streptomycin (Life Tech, #15070063). Cells were grown in dishes to a confluency of ~80-95%.

For H3K4me2 ChIP-seq, formaldehyde was added to media to a final concentration of 1% and incubated for 10 minutes on a shaker at room temperature. The reaction was quenched by adding glycine to a final concentration of 125 m*M* followed by 5 minutes of incubation on a shaker at room temperature. The cells were collected by centrifugation at 500 x g for 5 min at 4 °C and washed twice with ice-cold PBS. The cell pellet was resuspended in 10 ml ice-cold cell lysis buffer (5 mM HEPES (pH8), 85 m*M* KCl, 0.5% NP-40) with protease inhibitors (cOmplete^™^ ULTRA Tablets, Mini, EDTA-free, EASYpack Protease Inhibitor Cocktail, Roche # 05892791001) and incubated for 10 minutes at 4°C. Nuclei were released by 10 strokes with a Wheaton Dounce Homogenizer (tight pestle). The crude nuclear extract was collected by centrifugation at 500 x g for 5 min at 4°C, resuspended in 1 ml ice-cold nuclear lysis buffer (50 m*M* HEPES (pH 8), 10 m*M* EDTA, 0.5% N-Lauroylsarcosine with protease inhibitors) and incubated for 20 minutes at 4°C. After addition of 1 ml nuclear lysis buffer samples were sonicated using a Diagenode Bioruptor for 18 cycles (30” ON / 30” OFF) on “high”. After sonication samples were centrifuged at 14000 x g for 10 minutes at 4°C and the supernatant was aliquoted to DNA-low binding tubes. The chromatin was flash frozen in liquid nitrogen and stored at −80°C.

Protein A Sepharose (PAS) beads (Sigma #P9424) were washed twice with RIPA140 (140 m*M* NaCl, 10 mM Tris-HCl (pH 8), 1m*M* EDTA, 1% Triton X100, 0.1% SDS, 0.1% Na-Deoxycholate) with proteinase inhibitors and 1mg/ml BSA (Sigma #A7906) and incubated overnight. Chromatin was thawed on ice. 50μg of Chromatin was used per ChIP experiment. RIPA140 with proteinase inhibitor was added to a total volume of 1 ml and incubated with 2 μg of H3K4me2 antibody (Abcam, ab32356, Lot#: GR209821-1; or Epicypher 13-0013, Lot#: 14247001) for 16 hours at 4°C on a rotating mixer (40 rpm). 1% of the chromatin was used as input controls. Blocked beads were added to the chromatin-antibody complex solution and incubated for 3 hours at 4°C on a rotating mixer at 40 rpm. Complexes were washed once with 1ml RIPA140, 4 times with 1 ml RIPA500 (500 m*M* NaCl, 10 m*M* Tris-HCl (pH 8), 1m*M* EDTA, 1% Triton X100, 0.1% SDS, 0.1% Na-Deoxycholate) for 10 minutes each. Complexes were subsequently washed once in 1 ml LiCl-Buffer (250 mM LiCl, 10 mM Tris-HCl (pH 8), 1mM EDTA, 0.5% NP-40, 0.5% Na-Deoxycholate) and TE (10mM Tris-HCl (ph 8), 1m*M* EDTA) for 2 minutes each. Between each wash, beads were spun down at 500 x g for 2 minutes and the supernatant was discarded. Beads were resuspended in 100 μl TE and RNase A was added to a final concentration of 50 μg/ml followed by incubation at 37°C for 30 minutes. The samples were adjusted to a final concentration of 0.5% SDS and Proteinase K was added to a final concentration of 500μg/ml. Proteins were digested at 37°C for 90 minutes followed by reverse cross-linking overnight at 65°C. DNA was purified using phenol-chloroform extraction followed by ethanol precipitation. Libraries were prepared using the NEXTflex qRNA-Seq Kit v2 from Bioo Scientific (Catalog #5130–11) and sequenced on an Illumina NextSeq 500 sequencer.

For ATAC-seq, 200,000 cells were subjected to tagmentation as described (Buenrostro et al., 2013) with a total of 13 or 15 PCR cycles.

For PEAT data, *D. melanogaster* S2 cells were grown to a density of 3 million per ml in Schneider’s Medium (Invitrogen) supplemented with 10% FCS and 1x Antibiotics. PEAT library was constructed as described ^28^.

### Previously published datasets

ChIP-seq datasets for H3K4me1, H3K4me2, H3K4me3, and H3K27ac from *C. elegans* whole l3 stage were downloaded from data.modencode.org ^49^ corresponding to experiment IDs 5048, 5157, 3576, and 5054, respectively. ChIP-seq datasets for H3K4me1, H3K4me3, and H3K27ac from *D. melanogaster* S2 cells were downloaded from the Gene Expression Omnibus (GEO; Series GSE41440;^36^). ENCODE ChIP-seq datasets for H3K4me1, H3K4me2, H3K4me3, and H3K27ac from *H. sapiens* GM12878 cells were downloaded corresponding to GEO sample IDs GSM733772, GSM733769, GSM945188, and GSM733771, respectively, as well as input control sample IDs GSM733742 and GSM945259.

GRO-cap datasets for *H. sapiens* GM12878 cells (Series GSE60456;^12^) and *C. elegans* whole L3 stage (Series GSE43087;^4^) were downloaded from GEO.

ATAC-seq datasets for *H. sapiens* GM12878 cells were downloaded from GEO (Series GSE47753;^25^).

### Data processing

G(P)RO-cap datasets were subjected to adapter removal using cutadapt ^50^ as was ATAC-seq using flexbar ^51^ prior to mapping. Reads were then mapped with Bowtie2 ^52^ with default settings, including the parameter _X 1500 for paired-end datasets, to the hg19, ce6, or dm6 genome assemblies, followed by removal of multi-mapped reads from the resulting .sam files. ChIP-Seq data sets were aligned using bowtie2 with default parameters and reads that had more than 2 mismatches and did not align uniquely were removed. The sequencing library for S2 H3K4me2 ChIP-Seq data set was prepared with Unique Molecular Identifiers, therefore the first 9 bases of each read were removed using flexbarv2.4 ^51^ and reads were aligned to dm6 genome build using bowtie2 in paired-end mode keeping only concordantly aligned mates. All ChIP-seq datasets were collapsed using samtools rmdup ^53^ and ATAC-seq datasets were collapsed using MarkDuplicates.jar from Picard tools (http://broadinstitute.github.io/picard). Duplicates were not removed from P/GRO-cap datasets. ATAC-seq read pairs with fragments greater than 50bp were kept for further processing. Start sites of ATAC-seq reads were extended by 15 basepairs upstream and 22 basepairs downstream in a stranded manner, to account for steric hindrance of the transposition reaction ^54^. All reads that intersected ENCODE blacklisted regions (https://sites.google.com/site/anshulkundaje/projects/blacklists) were removed and all replicate BED files were concatenated together for peak calling and signal generation. Signal bigwig files for ATAC-Seq were generated using JAMM signal generator pipeline ^26^.

PEAT data was processed as follows. Fastq files from each mate were first matched and trimmed for the 5’end adapter using cutadapt ^50^ (parameters -a GTTGGACTCGAGCGTACATCGTTAGAAGCT -O 30 -m 20 --untrimmed-output). The sequences that were not matched for 5’end were then matched and trimmed for the 3’end adapter using cutadapt ^50^(parameters -a GTCGGATAGGCCGTCTTCAGCCGCCTCAAG -O 30 -m 20 --untrimmed-output). The two resulting fastq files matching each end were combined, reverse complemented, and then unpaired mates discarded and paired mates matched based on read IDs using custom scripts. The resulting paired fastq files were then mapped using STAR ^55^ (parameters--outFilterMultimapNmax 1 -- outFilterMismatchNmax 1--chimSegmentMin 30--chimJunctionOverhangMin 30--outFilterIntronMotifs RemoveNoncanonicalUnannotated--alignIntronMin 20--alignIntronMax 1000000--alignEndsType EndToEnd--alignMatesGapMax 1000000--alignSJoverhangMin 12--alignSJDBoverhangMin 3) to the dm6 genome assembly.

### ATAC-seq peak calling, annotation and selection

ATAC-seq peaks were called using JAMM v1.0.7rev5 setting the bin size to 100 and _e to “auto” for the gm12878 data set. The “all” list output from JAMM was used for gm12878 data set and the “filtered” output was used for the two other data sets. Only peaks that are larger than 50 basepairs were kept. To ensure that G(P)RO-cap start site counts in promoter regions are not underestimated due to narrow-width peak calls, final peaks were extended by 75 basepairs in each direction and overlapping peaks were merged. Extended, merged, ATAC-seq peaks were annotated as promoters if they were within +/−200bp from Gencode defined transcript starts for *H. sapiens*, +/−400bp from flybase defined transcript starts for *D. melanogaster*, and +/−500bp from refGene defined gene starts for *C. elegans*. Peaks that fell within these distances on both strands were considered known bidirectional promoters and removed from the analysis. Peaks were annotated as intergenic if they were not annotated as a promoter and did not intersect known transcript boundaries from the same databases.

ATAC-peaks containing confident transcription start sites from nascent RNA datasets were selected based on empirical distributions of read 5’ends per base from control regions. Control regions were selected as follows: first all ATAC-seq peaks containing at least one nascent RNA read 5’end on at least one strand were selected, then windows equal in size to a given ATAC-seq peak were taken immediately downstream of that ATAC-seq peak on both strands (i.e. higher coordinates on the plus strand and lower coordinates on the - strand), and any overlapping regions with other peaks were subtracted out. These windows were taken to represent frequently observed signal background within gene bodies that is likely to come from technical issues in the GRO protocols (i.e. non-nascent RNA contamination or inefficient cap-selection) and therefore are unlikely to represent true transcription start sites. The empirical distribution of nascent RNA read 5’ends per base was constructed across all the bases in the control windows and cutoffs were determined using the 0.999 quantile for *C. elegans* (17 read 5’ends) and *D. melanogaster* (8 read 5’ends), and the 0.9999 quantile for *H. sapiens* (14 read 5’ends). Thus, we had two sets of ATAC-seq peaks for downstream analysis: Set A contained at least one base with these numbers of reads on the forward strand only and Set B contained at least one base with these numbers of reads on both the forward and reverse strands. Set B peaks were filtered for convergent TSS pairs defined when the bases on both strands containing the most read 5’ends are both over the determined cutoff and are situated downstream from each other.

Non-extended peaks called using JAMM setting -m narrow were used for promoter peak width and distance between TSS and peak edge analysis (supplementary Figure 3A and B). Distance of TSS to the ATAC-Seq peak edge was determined using TSSs defined via extended, merged ATAC-Seq peaks (see above) and narrow peak edges, allowing for TSS occurring outside, downstream of the peak (negative distances) and instide the peak (positive distances). Peak width and distance from TSS to peak edge were done using the forward TSS of both Set A and Set B promoter peaks (see above). If the absolute TSS distance to the edge was larger than 200bp, it was not included in the boxplot.

HeLa-S3 DNase-I peaks and 5’GRO-seq data used in Figure S1A were obtained from Duttke et al. 2015b,

### Histone Modification Peak Calling and Signal Files

All histone modification peaks were called using JAMM v1.0.7rev5 setting the bin size to 150 and -r to “window” ^26^. For S2 H3K4me2 data set, peaks were called using JAMM v1.0.7rev5 in paired-end mode. The “filtered” output peaks produced by JAMM was used.

Histone modification bigwig signal tracks were generated using deepTools ^56^ bamCoverage at 10bp resolution using the fragment length obtained by JAMM and setting normalization to RPKM. To generate average meta-plots, deepTools computeMatrix was used at single basepair resolution. The bigwig files were also used to define the features for partial correlations and the predictive linear model.

### TSS sequence model

The TSS sequence model initially described by Frith and colleagues ^33^ was used as described previously ^11^. Set A ATAC-Seq peaks (see above) were used for model training as follows: a window +/−50 bp surrounding the position with the most nascent RNA read 5’end counts within the ATAC-seq peak was used to train the TSS sequence model. The model was then run either on the same corresponding windows from both strands of the selected promoters (Set B ATAC-Seq peaks, Figure 2B). Midpoints between forward and reverse TSSs served as negative controls for sequence model scores. Alternatively, the model was run on windows surrounding all bases on each strand that had at least one nascent RNA read 5’end and the scores summed per strand for the partial correlation analysis (Figure 2C; Figure 4).

### TSS distribution pattern score

Distribution pattern (*DP*) scores were calculated based on the equation for Shannon entropy similar to a previously described method ^32^. Specifically,

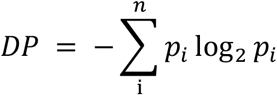

where *p* is the probability of a nascent RNA read 5’end at position *i* for a given strand of an ATAC-seq peak and *n* is all the positions for that strand that have at least one read 5’end.

### GC content and skew

GC percentage was calculated in a sliding 50 bp window with a step size of 1 bp along a given region and then taking the positional mean across all selected regions. GC skew was calculated as follows:

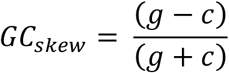

where *g* and *c* are the number of G and C nucleotides in a 50 bp window slid along a given region with a step size of 1 bp. Positional means were then calculated across all regions.

### Mixture modeling

Mixture modeling of ATAC-seq peak nascent RNA read 5’end count ratios was performed using the R package Mclust with default parameters ^57^.

### Chromatin State Hidden Markov Model

Histone modification peaks were processed for chromatin state HMM as previously described ^11^. Chromatin states were then obtained as previously described ^11^ at 10 basepair resolution using multivariate normal distribution for the emission probabilities but with the following changes: only sequences that were at least 500 basepairs long were kept for Baum-Welch training which was done setting the transition probability matrix to 0.9 at the diagonal and 0.1 / (n – 1) at all other entries where n is the number of states, and segmentation was done using posterior decoding. Baum-Welch was run on chromosome 1 for gm12878 and on all chromosomes for S2 and L3. Scripts for baum-welch training and posterior decoding are available at https://github.com/mahmoudibrahim/hmmForChromatin. The two H3K4me1-only states were summed and plotted as one line in state coverage plots in all figures.

### Partial Correlation Analysis

Histone modification features were defined as the maximum histone modification ChIP-Seq signal in a 1kb window downstream and upstream of the non-extended ATAC-seq peaks (see above) for forward and reverse directions respectively. Initiation rate features were the total cap-selected GRO-seq read 5’ends on each respective strand of the extended ATAC-seq peaks divided by ATAC-Seq peak width. Core promoter sequence features were the sum of all model scores for positions with at least one cap-selected GRO-seq read 5’end divided by ATAC-Seq peak width (see above). ATAC-Seq features were defined as the maximum ATAC-Seq signal in the non-extended ATAC-seq peaks.

Spearman partial correlation coefficient were then obtained using the R package ppcor ^58^ and heatmaps were plotted using the R package pheatmap ^59^.

### Transcription directionality model

Features were defined as in partial correlation analysis (see above). A linear mixture model was learned on all *H. sapiens* promoter regions that has confident forward and reverse initiation sites (see above) using the R package flexmix ^60^, setting the number of clusters to 2. This results in learning two linear models with distinct regression coefficients and assigning each data point a probability of belonging to each of the two mixture components. Each promoter region is then assigned to the mixture component that with the higher probability. To obtain predicted directionality ratios, the prediction from the mixture component that the promoter region belongs to is used.

For cross validation analysis, the data was split into 10 equal parts and model learning and clustering were repeated 10 times and each time the model predictive ability was tested on the held-out test set, summarized using the correlation coefficient. This was done separately for three different models, one that included both core promoter sequence score ratio and H3K4me3 ratio and two that included sequence score ratio only and H3K4me3 only.

### *D. melanogaster* Enhancer Analysis

STARR-Seq ^43^ peaks were obtained from the Stark lab website (http://www.starklab.org/data/arnold_science_2013/) and coordinates were lifted over to dm6 genome assembly. Peaks from both replicates were merged and extended by 200bp in each direction. For Figure 5C, promoters that belonged either Set A ATAC-Seq peaks or Set B peaks were chosen and stratified by whether their corresponding transcript intersected a STARR-Seq peak that intersected an ATAC-Seq peak. For Figure 5D, all promoter annotating ATAC-Seq peaks were stratified by whether they intersect a STARR-Seq peak.

## Author Contributions

S.A.L. and M.M.I. designed the project and performed the analysis with assistance from A.K. and advice from U.O.. S.A.L, M.M.I, and U.O. wrote the manuscript. A.K. and A.H. performed the ATAC-seq in *D. melanogaster* S2 cells. E.K. performed the ATAC-seq in *C. elegans* with advise from S.A.L., B.T., and A.H.. A.G. performed the *D. melanogaster* S2 cell H3K4me2 ChIP-seq with advice from R.Z. and A.H. prepared the library for sequencing. A.C. performed the *D. melanogaster* S2 cell PEAT.

## Acknowledgements

The authors would like to thank Hans-Hermann Wessels for help with S2 cell culture as well as James T Kadonaga, Sascha Duttke, Sven Heinz and Chris Benner for helpful comments and discussions.

---

**Figure S1.**
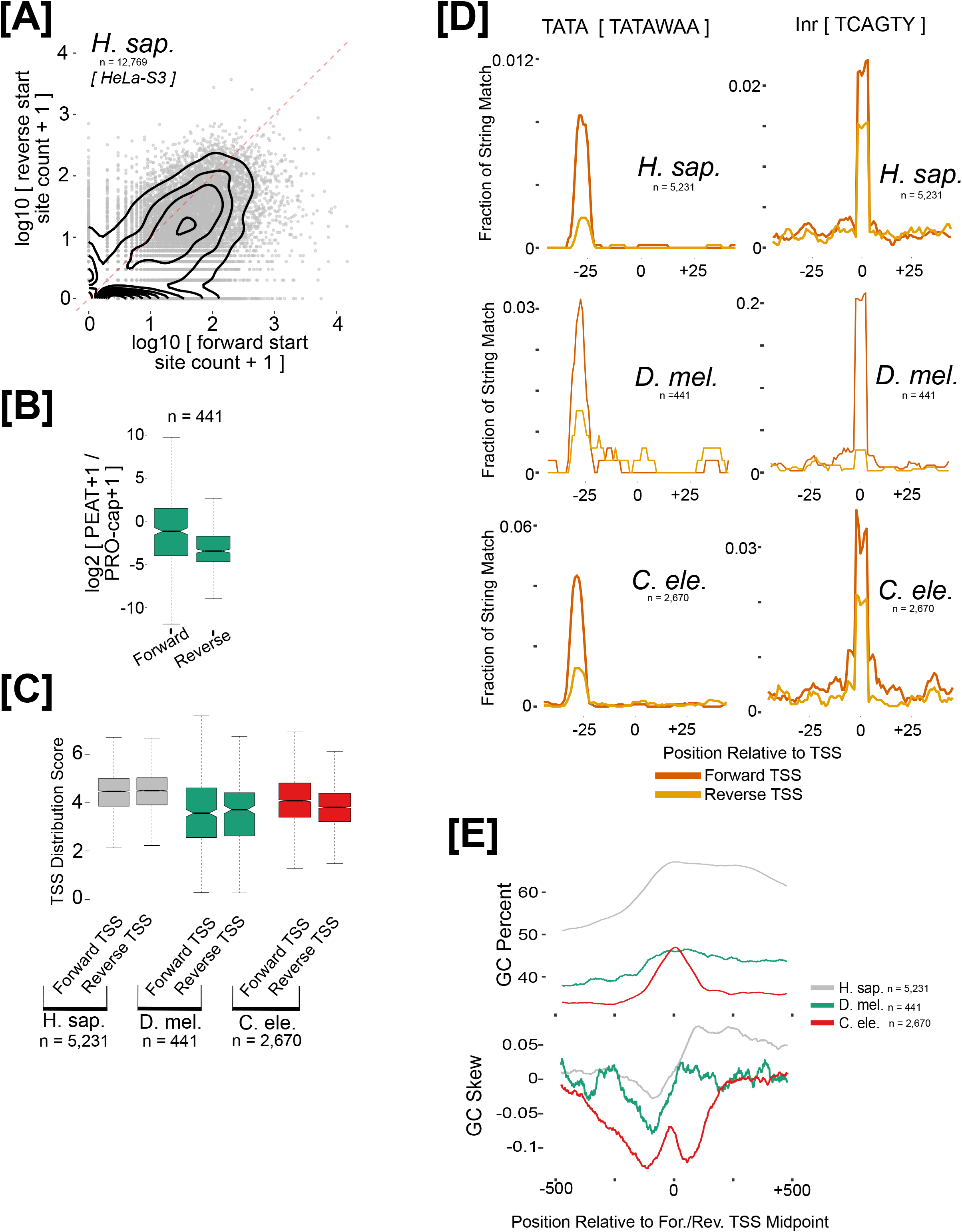
Promoter transcription directionality and divergent promoter characteristics. **A**. Forward versus reverse direction cap-selected GRO-seq counts plotted for promoter NDRs defined by DNaseI-seq in *H. sapiens* HeLa cells. **B**. Forward and reverse PEAT-to-PRO-cap count ratio distributions for *D. melanogaster* promoter NDRs containing significant forward and reverse TSSs. **C**. Forward and reverse TSS distribution entropy scores for promoter NDRs containing significant forward and reverse TSSs. **D**. Initiator and TATA-box consensus sequence string matches relative to forward and reverse TSS modes for promoter NDRs containing significant forward and reverse TSSs. **E**. Positional averaged analyses of GC percentage (top) and GC skew (bottom) in 50 bp sliding windows relative to midpoints between forward and reverse TSS modes for promoter NDRs containing significant forward and reverse TSSs.

**Figure S2.**
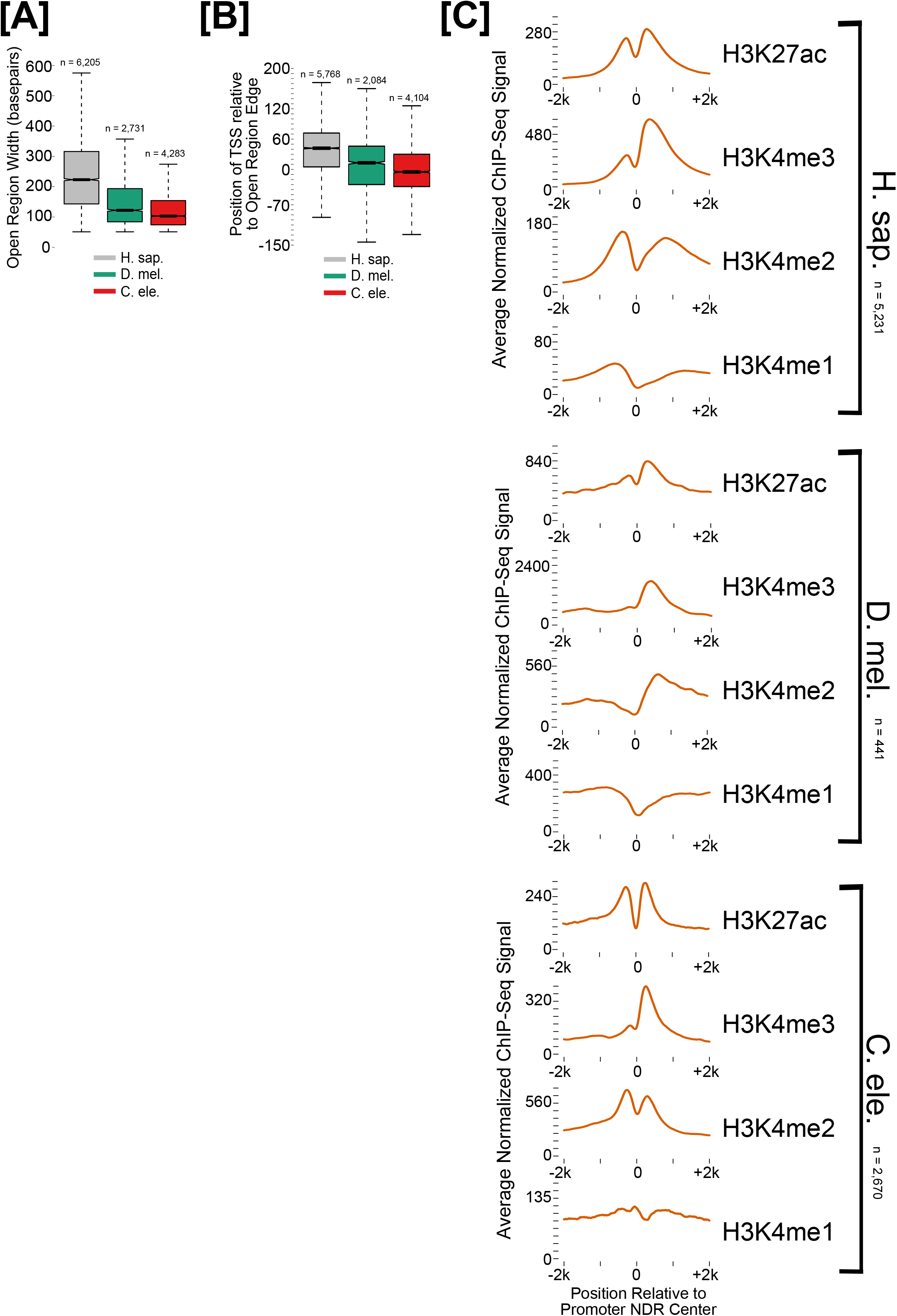
NDR width, TSS-edge distances, and histone PTM average analyses. **A**. Promoter NDR width distribution. **B**. Distributions of TSS positions relative to NDR edge (negative numbers indicate the TSS is downstream and outside of the NDR edge, positive number indicate the TSS is inside the NDR edge). **C**. Meta analyses of histone PTMs relative to midpoints of promoter NDRs containing significant forward and reverse TSSs.

**Figure S3.**
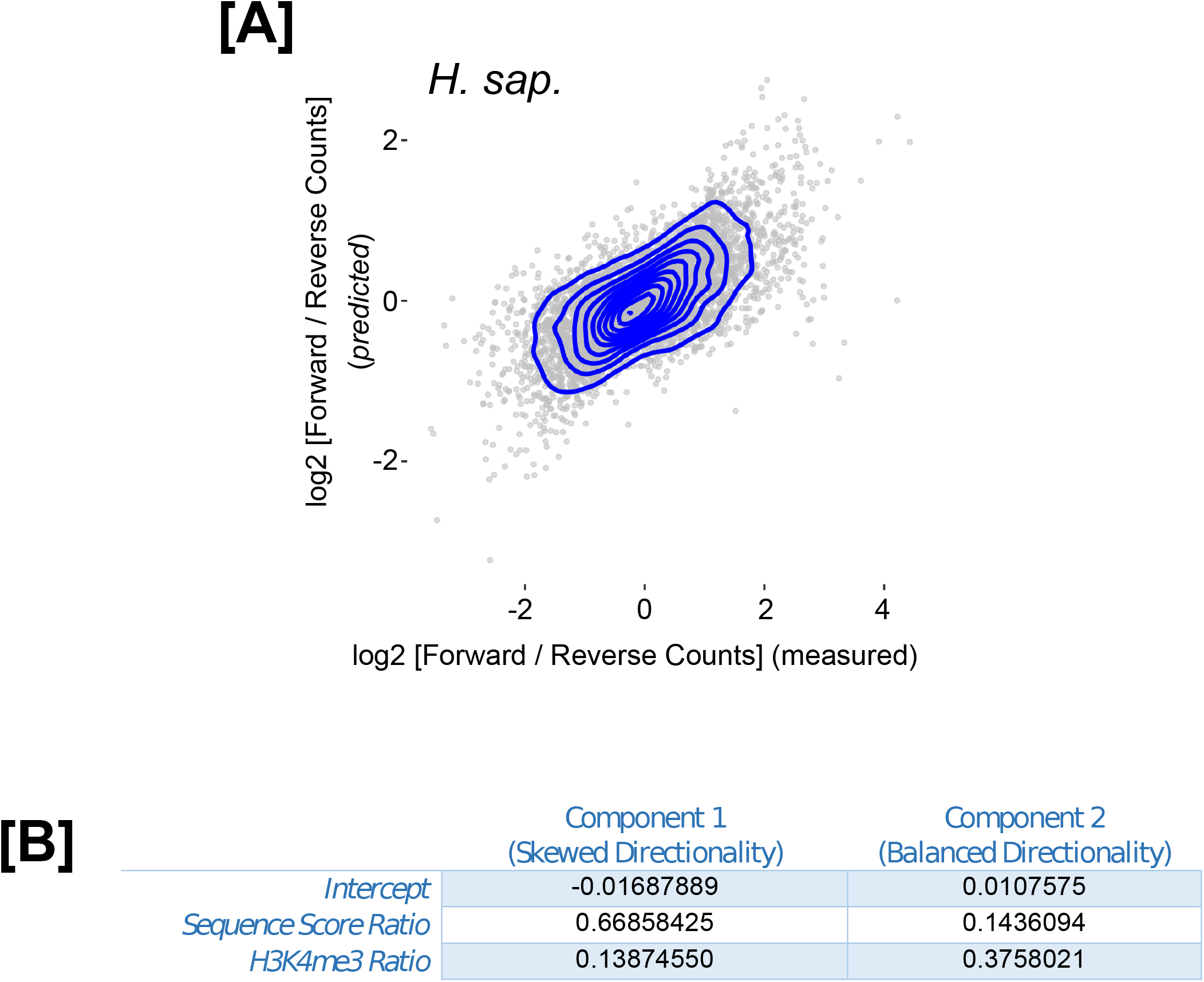
Clustering of promoter regions based on sequence and chromatin features. **A**. Same as Figure 4A but with data points belonging to each linear regression model plotted together. **B**. Regression coefficients learned when training the model on all promoter regions (same model as in Fig. 4A).

**Figure S4.**
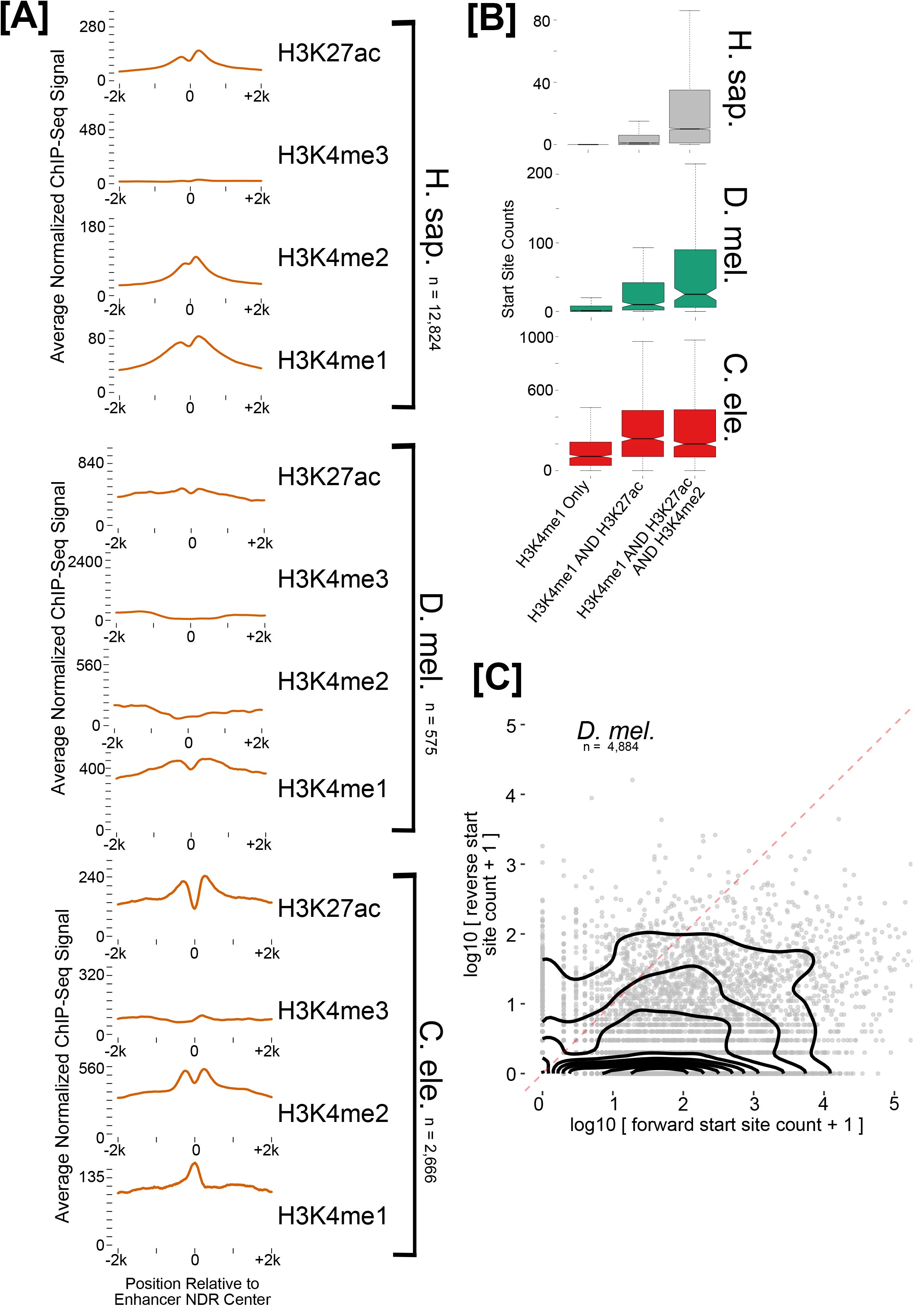
Distal NDR characteristics and *D. melanogaster* enhancers. **A**. Meta analyses of histone PTMs relative to midpoints of intergenic NDRs intersecting at least H3K27ac and H3K4me1 peaks. **B**. Total cap-selected GRO-seq count distributions for distal NDRs intersecting H3K4me1 only (n = 12,408(Hs) 802(Dm) 484(Ce)), H3K4me1 and H3K27ac only (n = 1,002(Hs) 284(Dm) 722(Ce)), or H3K4me1, H3K4me2 and H3K27ac (n = 7,324(Hs) 225(Dm) 741(Ce)). **C**. Forward versus reverse direction cap-selected PRO-seq counts plotted for *D. melanogaster* promoter NDRs not intersecting STARR-seq peaks.

**Table S1.** Full partial correlation table between total cap-selected GRO-seq counts (Expr.), ATAC-seq signal, TSS distribution entropy (Dist.) and core promoter sequence score sums (Seq.) in forward and reverse directions for promoter NDRs with significant forward and reverse TSSs

**Table S2.** Full partial correlation table for active promoter histone PTMs and ATAC-Seq in forward and reverse directions for promoter NDRs with significant forward and reverse TSSs.

